# Limb-on-a-Chip: An All-Hydrogel Platform for Scalable and Reproducible Engineering of Neuromuscular Tissues

**DOI:** 10.1101/2025.10.12.681910

**Authors:** Laura Schwendeman, Angel Bu, Sonika Kohli, Ferdows Afghah, Tamara Rossy, Ritu Raman

## Abstract

Diseases or injuries that impact neuromuscular tissues have a severe negative impact on human health, mobility, and quality-of-life, motivating the development of tissue engineered *in vitro* models of the motor control system. Current neuromuscular organoids and organ-on-a-chip platforms either rely on stochastic self-assembly that limits reproducibility or require complex microfabrication processes that preclude high-resolution imaging and scalable functional analysis. We have developed a Limb-on-a-Chip platform that addresses key challenges of current model systems by enabling reproducible and scalable manufacturing of neuromuscular tissues compartmentalized into “spinal cord” and “limb” chambers, while promoting biochemical crosstalk between cell types. Our fabrication method leverages 3D printed molds to perform 1-step micropatterning of an all-hydrogel chip containing precise features to guide muscle fiber alignment and motor neuron axonal outgrowth. We demonstrate the ability to co-culture motor neurons and skeletal muscles within this hydrogel platform, enabling tissue-wide readouts of muscle force as well as single cell-resolution measurements of muscle fiber calcium activity. Our accessible method for fabricating reproducible *in vitro* neuromuscular models that are compatible with high-resolution imaging and functional readouts provides a powerful new tool for investigating the neuromuscular interface in health and disease.

## INTRODUCTION

Pathologies that affect the neuromuscular system, namely skeletal muscles and the motor neurons that control their activity, have a severe negative impact on human health, mobility, and quality of life.^1^ Neuromuscular pathologies impact people across the age spectrum and can stem from a variety of causes, ranging from inherited muscular dystrophies to traumatic injuries to neurodegenerative diseases.^2,3^ *In vitro* models of the neuromuscular interface have emerged as useful tools for investigating the underlying mechanisms of disease onset and progression, as well as to develop new therapeutic approaches that preserve or restore function.^4–6^

They key functional unit of the neuromuscular interface is the motor unit (a motor neuron and the downstream muscle fibers it controls), and several studies have established the ability to generate functional motor units *in vitro*. Among the most well-established approaches for generating motor units *in vitro* are protocols that rely on self-assembly.^7^ For example, several studies have demonstrated the ability to form functional neuromuscular junctions within 3D organoids derived from stem cells, including both self-organizing neuromuscular organoids^8,9^ and “assembloids” of motor neuron spheroids fused with skeletal muscle spheroids.^10,11^ While these approaches enable forming multiple independent and functional motor units within a mm-scale tissue, the large size of these tissues (∼2-5 mm diameter) limits compatibility with high-throughput imaging of neuromuscular junction architecture or function.^12^ Moreover, these tissues are not compatible with functional readouts of contractile force. To combat this drawback, researchers have demonstrated that 2D co-cultures of skeletal muscle and motor neurons enable readily visualizing neuromuscular junction formation and activation.^12–15^ However, delamination of muscle monolayers from underlying substrates makes it difficult to preserve such 2D cultures over several weeks in culture, and often preclude functional readouts of contractile force.^16–19^ Overall, the stochastic processes that drive formation and maturation of both the 2D and 3D systems described above make it difficult to reliably place, stimulate, and quantitatively monitor independent motor units within *in vitro* self-assembled neuromuscular models.^20^

By contrast, organ-on-a-chip approaches for engineering neuromuscular tissues enable fabricating systems with more reproducible architectures. These techniques typically rely on multi-step microfabrication processes to generate “chips” composed of precisely defined physical compartments that house 3D motor neuron spheroids and 3D muscle tissues, including architectural features that promote formation of anisotropically aligned muscle fibers that mimic native tissue morphology.^20–22^ More complex models even separate neurons and muscles into separate compartments connected by microchannels for axons to grow through, enabling stimulation of motor neurons using chemical or optogenetic techniques to trigger downstream muscle contraction.^23,24^ Importantly, both non-compartmentalized and compartmentalized organ-on-a-chip models couple 3D muscle tissues to flexible cantilevers, enabling quantitative readouts of force.^25^ While such organ-on-a-chip platforms thus improve reproducibility of tissue architecture, their reliance on 3D tissues limits the ability to precisely visualize individual motor units, or quantify the activity of individual muscle fibers, similar to 3D neuromuscular organoids. The current state-of-the-art^26^ thus motivates developing an *in vitro* platform that combines the precise tissue patterning and force readouts of 3D organs-on-a-chip with the ease of high-resolution imaging offered by 2D self-organizing co-cultures.

We have developed a 2D compartmentalized platform that addresses this challenge and enables scalable and reproducible engineering of the neuromuscular interface. Our device design takes inspiration from the native spinal cord in humans and other mammals. *In vivo*, motor neuron soma are housed within the spinal cord, and project axons outwards to innervate skeletal muscle tissue in the periphery, with multiple motor units working together to control a downstream limb (**Figure 1**). Likewise, in our spinal cord inspired-device, motor neuron soma are isolated in separate compartments, and connected via micro-channels to a downstream mm-scale 2D muscle tissue. The fabricated tissue thus represents a Limb-on-a-Chip, composed of multiple independent units. A key advantage of our platform is that it uses precision engineering to compartmentalize motor neurons and skeletal muscle, mimicking the reproducible architecture of 3D organs-on-a-chip, while retaining a 2D morphology at the neuromuscular interface, similar to self-organized 2D co-cultures. Additionally, the open-well format of our device enables co-culturing muscles and neurons in the same media, in contrast to microfluidic devices that fluidically separate the two cell types. Promoting biochemical crosstalk between muscles and motor neurons is known to promote tissue maturation, highlighting the benefit of leveraging an open-well format.^27,28^ Furthermore, unlike traditional organ-on-a-chip devices, which rely on multi-step microfabrication protocols to fabricate complex microfluidic structures, our Limb-on-a-Chip relies on a 1-step hydrogel molding protocol that leverages a 3D printed stamp to precisely and rapidly pattern an all-hydrogel device. In this study, we first present and validate the reproducibility of our Limb-on-a-Chip fabrication protocol, and demonstrate the ability to co-culture contractile muscles and motor neurons in separate compartments. We anticipate that our accessible protocol will enable researchers to fabricate reproducible, scalable, and functional models of the neuromuscular interface, thus advancing the state-of-the-art in neuromuscular disease modeling.

**Figure 1.**
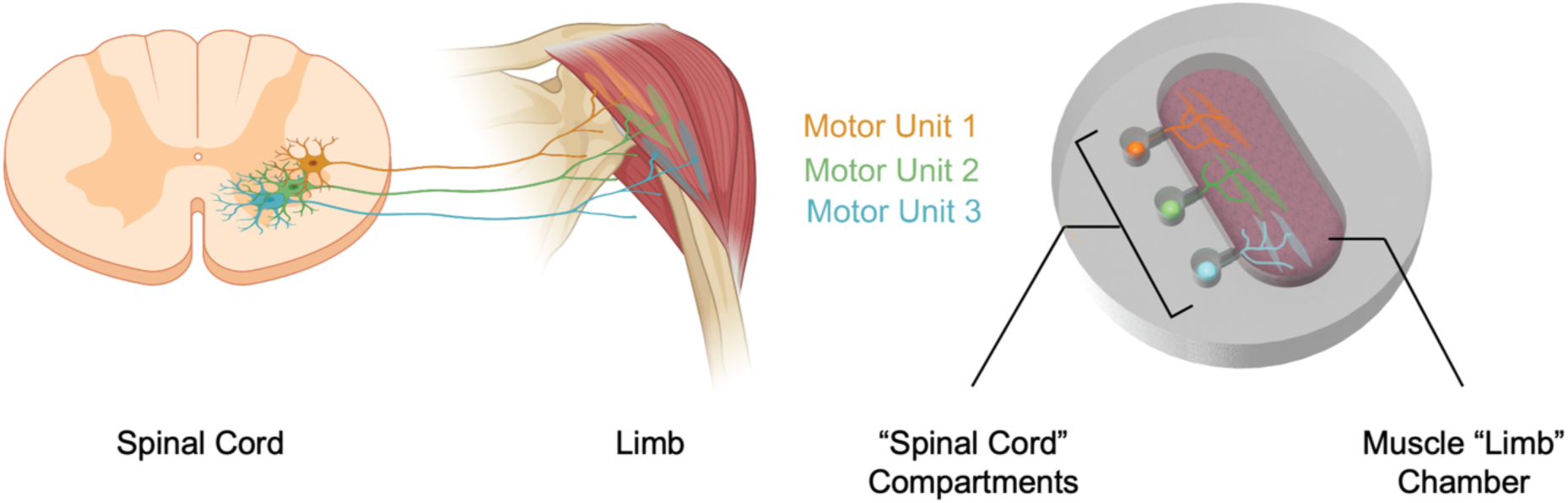
Bioinspired Limb-on-a-Chip Design. Compartmentalization of motor neurons in the spinal cord enables forming independently controllable motor units in a downstream limb. *In vivo* architecture (left) is mimicked in our *in vitro* Limb-on-a-Chip device (right).

## RESULTS

### Limb-on-a-Chip Design and Fabrication

Reproducible fabrication of functional neuromuscular interfaces in 2D requires methods to anisotropically align and mature contractile skeletal muscle monolayers. We have recently developed a technique termed STAMP (simple templating of actuators via micro-topographical patterning)^16^ that leverages a 3D printed mold to pattern micro-scale grooves in a hydrogel substrate, thus providing a physical template to guide fusion of myoblasts into a monolayer of precisely aligned multinucleated muscle fibers. While our previous study demonstrated that STAMP enables reproducible fabrication of muscles in monoculture, adapting STAMP for 2D neuromuscular organ-on-a-chip devices requires a method to precisely compartmentalize muscles and motor neurons in different chambers, while integrating patterning features that both promote alignment of muscle fibers and guide axonal outgrowth from motor neurons.

We designed a Limb-on-a-Chip STAMP, composed of a mm-scale muscle “limb” compartment containing aligned parallel 25 μm width grooves (dimensions previously demonstrated to promote efficient muscle fiber alignment^16^) connected to three separate motor neuron 750 μm diameter compartments (mimicking separate motor units within a spinal cord) via 500 μm long micro-channels (**Figure 2a-b**). Chip design and dimensions were chosen to ensure clear visual separation of muscle and neuron compartments, enabling seeding myoblasts and motor neurons into segregated chambers. Since our motor neuron spheroids are ∼500 μm in diameter on average,^21,27^ while individual myoblasts are ∼13 μm in diameter (Figure S1), micro-channels connecting the muscle and motor neuron compartments were designed to be 10 μm wide at the edge of the muscle chamber to prevent myoblast infiltration into neuron compartments after compartmentalized cell seeding. Preliminary experiments showed that STAMPs with micro-channels that preserved a 10 μm width throughout the channel length tended to deform due to buckling of the thin long feature, thus limiting the ability to fabricate precise axon guidance channels within hydrogels (Figure S2a). We thus leveraged a tapered channel design that transitioned from a 50 μm width at the neuron chamber interface to a 10 μm width at the muscle chamber interface, enabling preserving channel shape fidelity in both the STAMP and the molded hydrogel without buckling (Figure S2b).

**Figure 2.**
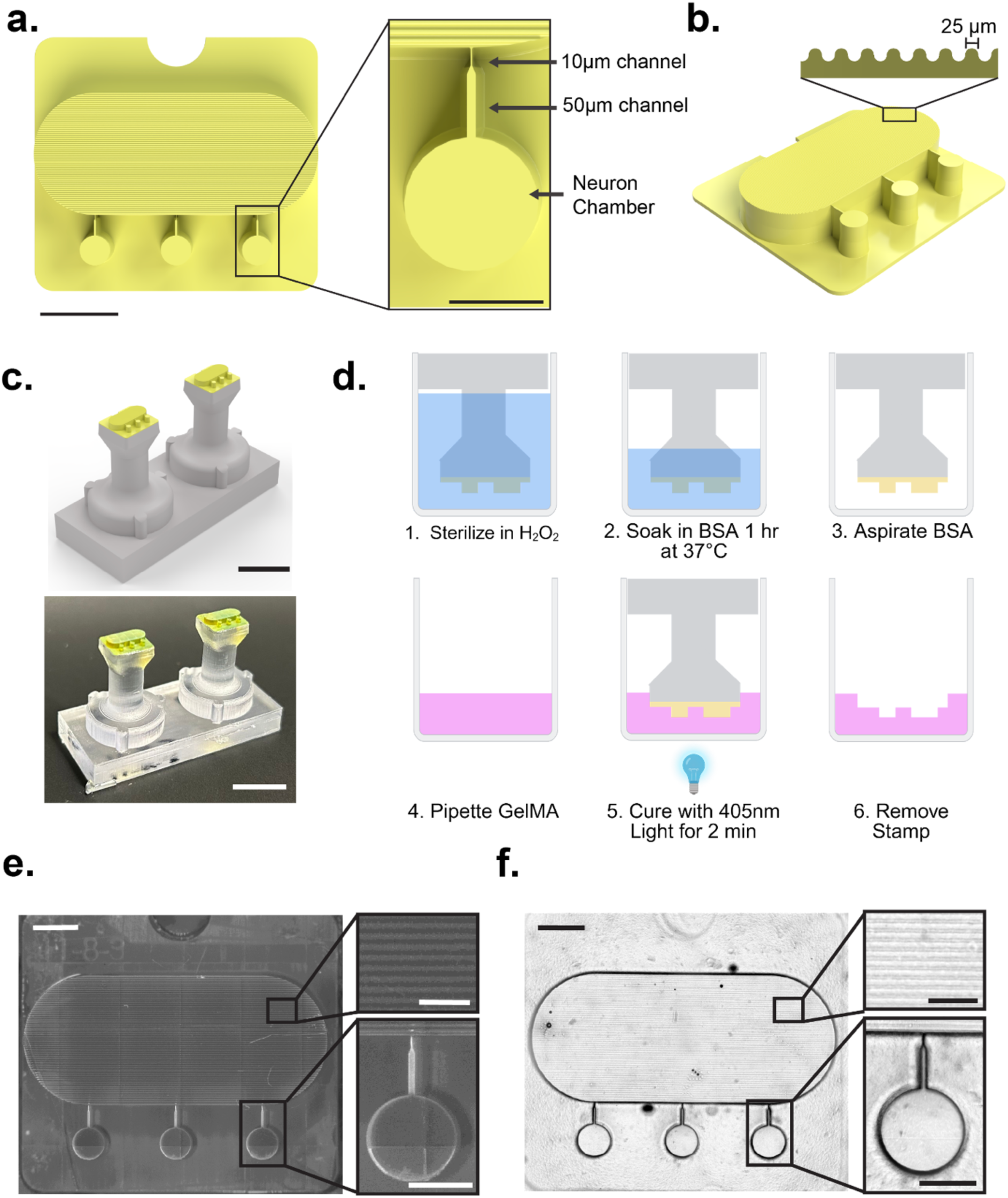
Limb-on-a-Chip Design and Fabrication. **a)** Top-down CAD rendering of Limb-on-a-Chip STAMP mold (scale bar = 2mm). Inset: Top-down CAD rendering of neuron chamber region and connecting channel design consisting of a smaller 10 µm width channel region for preventing muscle infiltration followed by a reinforcing 50 µm width region that preserves channel integrity (scale bar = 500 µm). **b)** CAD rendering of Limb-on-a-Chip STAMP highlighting the 25 µm grooved features of the muscle chamber region. **c)** Comparison of CAD rendering of assembled STAMP and holders (top) to the actual physical device (bottom). Holders enable precisely aligning STAMPs within 24-well plates. (scale bars = 1 cm). **d)** Overview of process used to pattern 7.5% GelMA Limb-on-a-Chip hydrogels with STAMPs. **e)** Brightfield image of representative Limb-on-a-Chip STAMP surface (scale bar right = 1 mm; scale bars left = 500 µm). **f)** Confocal transmitted channel image of representative casted Limb-on-a-Chip hydrogel (scale bar right = 1 mm; scale bars left = 500 µm).

Fabricated STAMPs were mounted onto a 3D printed holder (Figure 2c), enabling easy handling during the hydrogel chip molding process. STAMP-holder assemblies were sterilized, coated with bovine serum albumin (BSA) to prevent hydrogel adhesion during molding, and used as a template to pattern gelatin methacrylate (GelMA) hydrogels into the desired Limb-on-a-Chip architecture (Figure 2d). Imaging of STAMPs and STAMPed hydrogels showed that the compartmentalized chip architecture, as well as the microscale design features within each chamber, were accurately transferred from the mold to the all-hydrogel chip (Figure 2e-f).

### Limb-on-a-Chip Design Validation

Compartmentalized seeding and differentiation of motor neurons and skeletal muscles within separate chambers of the Limb-on-a-Chip first required validating that fabricated devices maintained their precisely patterned geometries over long-term culture in liquid media. In our previous proof-of-concept study of muscle monocultures, we showed that STAMP could be used to pattern the topography of fibrin hydrogels, but that swelling and remodeling of the soft fibrin (0.05 ± 0.009 kPa storage modulus at 10 rad/s, Figure S3b) over several days in culture reduced shape fidelity of the patterned hydrogel over time,^16^ precluding the ability to fabricate compartmentalized 2D co-cultures. To address this challenge, we fabricated Limb-on-Chip devices from stiffer GelMA hydrogels (0.3 ± 0.016 kPa storage modulus at 10 rad/s, Figure S3b) that still matched the mechanical properties of substrates typically used for muscle and motor neuron culture^27,29^ and then monitored the long-term fidelity of microscale features within the chip.

We obtained 3D confocal microscopy images of micro-grooves within the muscle chamber and used the open-source Segment Anything Model (SAM)^30^ to obtain a binary image of the channel features (Figure 3a). Comparing the measured hydrogel groove dimensions to the STAMP mold groove dimensions over 1 week showed that fabricated groove width closely matched the nominal value of 25 μm, and remained stable over 1 week while immersed in cell culture media (Figure 3b-e).

**Figure 3.**
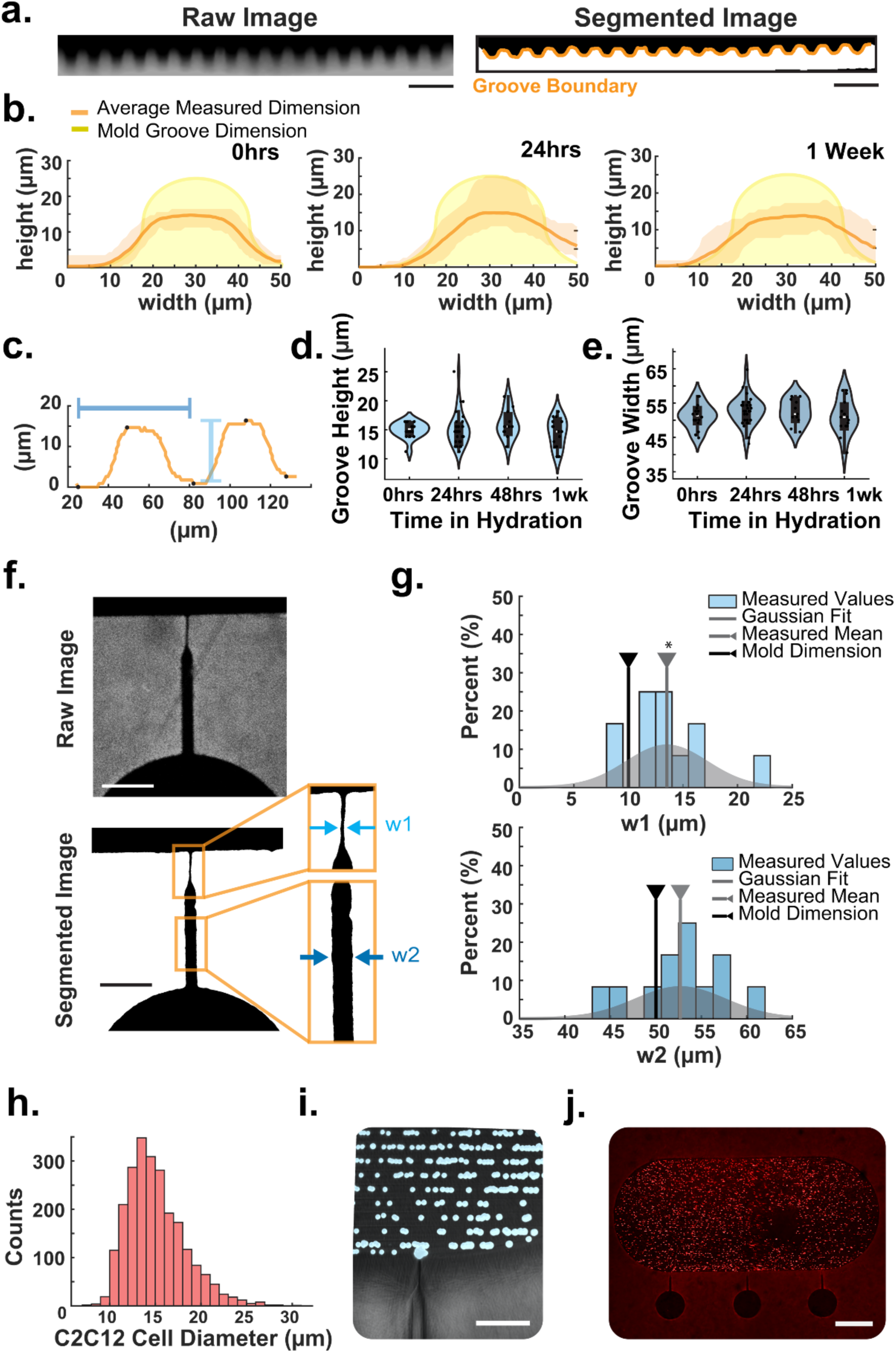
Limb on a Chip Design Validation. **a)** Right: Representative confocal z-stack cross section of hydrogel florescence in grooved regions of the Limb-on-a-Chip hydrogel muscle chamber region 0hrs after casting. Left: SAM segmented and binarized image of grooves (white) with a traced boundary used for data analysis shown in orange. (scale bars = 100 um). **b)** Average segmented groove dimensions (orange) from Limb-on-a-Chip hydrogels across 0hrs (left; n = 16), 24 hrs (middle; n = 29), and 1 week (right; n = 15) compared to the nominal 25 µm groove dimensions. Faded orange regions denote the bounds of the minimum and maximum measured dimensions at each timepoint. **c)** Representative groove traces from 0hrs after stamping (orange). Identified peak and valley locations are denoted with black dots. Each groove’s width is measured as the distance between subsequent valley locations (dark blue) and each groove’s height is measured as the distance between a valley point and the next following peak (light blue). **d)** Measured groove heights across multiple days after being hydrated in phosphate buffered saline (PBS). No statistical difference in measured groove height was found according to a Kruskal-Wallis test (H(3) = 1.8580; p = 0.6024); (0hrs: n = 16, 24hrs: n = 29, 48hrs: n = 11, 1wk: n = 15). **e)** Measured groove widths across multiple days after being hydrated in PBS. No statistical difference in measured groove width was found according to a one-way ANOVA (F(4) = 0.8585; p = 0.4722) (0hrs: n = 16, 24hrs: n = 29, 48hrs: n = 11, 1wk: n = 15). **f)** Representative confocal image of channel region in a Limb-on-a-Chip hydrogel (top) and the subsequent segmented binarized image. Two different regions of the channel were cropped to identify the region cast by the 10um region of the Limb-on-a-Chip STAMP mold (w1) and the 50 µm region of the channel (w2). **g)** Top: histogram of average measured w1 width across n = 12 channels (light blue) with a fit gaussian (gray; μ = 13.5171; σ = 3.676 µm) overlay. The average width (gray line) was found to be significantly greater than 10um according to a one sample t-test (t(11) = 3.3120; p = 0.0069). Bottom: histogram of average measured w2 width across n = 12 channels (dark blue) with a fit gaussian (gray; μ = 52.6909 um; σ = 4.9243 um) overlay. The average width (gray line) was not found to be significantly different from 50 µm according to a one sample t-test (t(11) = 1.893; p = 0.0850). **h)** Histogram of WT C2C12 diameters measured with Cellpose labeling of n = 2322 cells across 5 different brightfield images of myoblasts in suspension (mean = 15.08 µm; mode = 13.6713 µm; std. dev = 3.1547 µm). **i)** Representative confocal 20x objective image of fluorescent 10 µm polystyrene beads (red) seeded in the muscle chamber and staying separated from the neuron chamber (scale bar = 200 μm). **j)** Representative image of C2C12 muscle myoblasts labeled with a red fluorescent tag and seeded in the muscle chamber. Successful prevention of infiltration was observed at a 72.2% success rate across n = 18 channels. (scale bar = 1 mm).

Similarly, 3D confocal microscopy images of micro-channels separating the neuron and muscle chambers were segmented, measured, and compared to nominal STAMP dimensions (Figure 3f-g). While the wider 50 μm portion of the micro-channels closely matched mold dimensions, the narrow segment of the micro-channel was significantly wider than specified (13.5 μm ± 3.68 μm, as compared to mold dimensions of 10 μm), similar to the width of individual myoblasts (Figure 3h). Since average channel width may not be representative of the actual width at the muscle chamber interface, we conducted “leak testing” of channels with 10 μm fluorescent polystyrene beads and observed no visible infiltration of beads into channels (Figure 3i). These tests were further corroborated by using a pipette to load fluorescently labeled C2C12 mouse myoblasts into the muscle chamber of the open-well Limb-on-a-Chip (Figure 3j), demonstrated that myoblasts remained in the muscle chamber without infiltrating the micro-channels, enabling spatial segregation from the motor neuron compartment as desired.

### Differentiation and Visualization of Contractile Skeletal Muscle within Limb-on-a-Chip

Functional *in vitro* neuromuscular models require the ability to monitor and quantify skeletal muscle contractile force. We have previously shown that C2C12 mouse myoblasts seeded on micro-patterned fibrin hydrogels containing 25 μm grooves efficiently fuse into globally aligned multinucleated muscle fibers.^16,31^ To validate compatibility of our established protocols^32^ with GelMA hydrogels, we seeded wild-type myoblasts (WT C2C12) into the muscle compartment of a Limb-on-a-Chip. Myoblasts were maintained in growth media containing fetal bovine serum (FBS) until confluence, and then transitioned to differentiation media containing horse serum (HS) and human insulin-like growth factor-1 (IGF-1) to promote fusion into mature muscle fibers (Figure 4a). We concurrently seeded and differentiated C2C12 myoblasts engineered to express a genetically encoded calcium sensor (RGECO C2C12) in Limb-on-a-Chip devices using the same protocol. Calcium sensors enable visualizing depolarization of individual muscle fibers within a tissue (Video S1), thus enabling single cell-resolution in visualizing fiber activation, in addition to tissue-wide functional force readouts.

**Figure 4.**
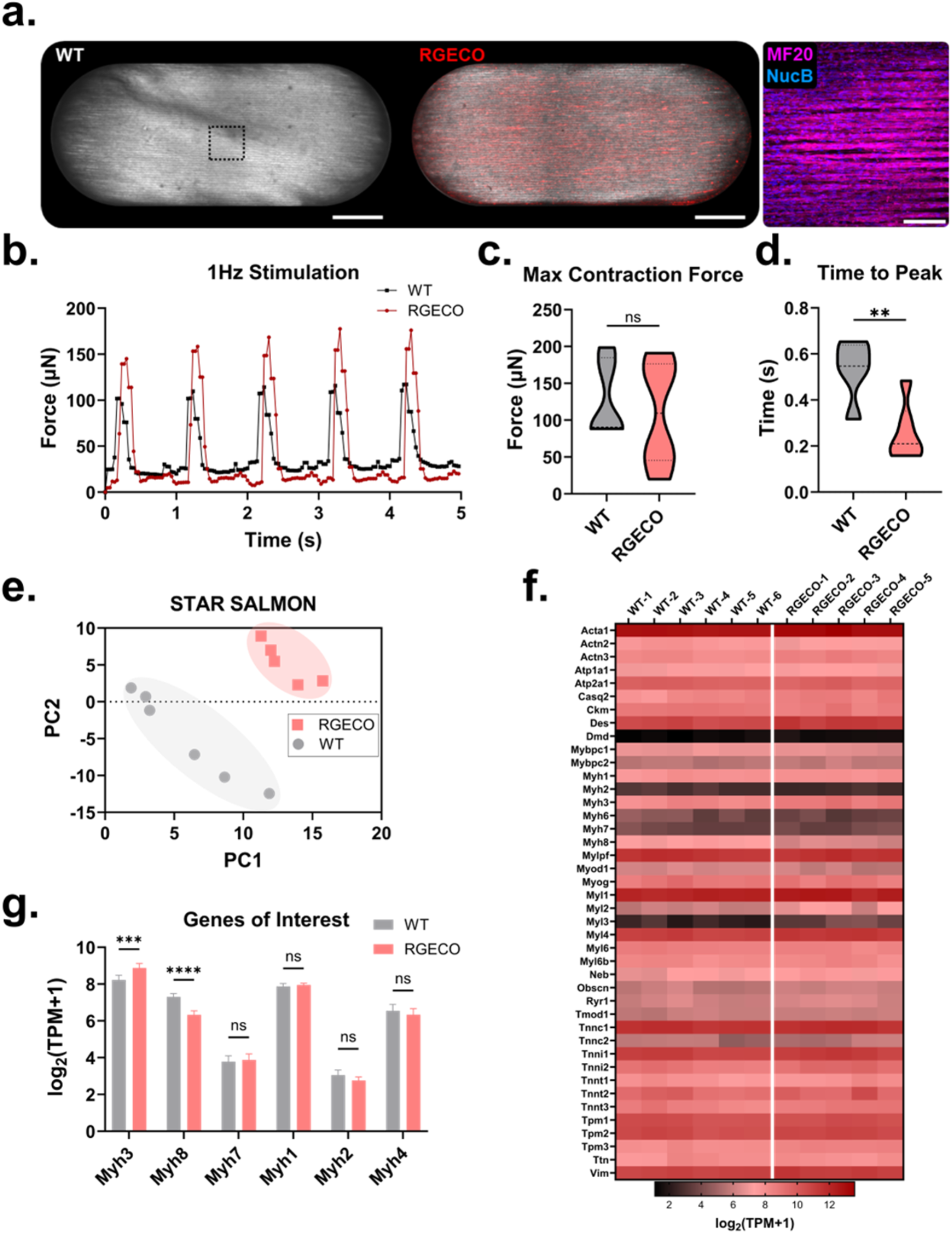
Differentiation and Visualization of Contractile Skeletal Muscle within Limb-on-a-Chip. **a)** Representative images for mature WT and RGECO C2C12-derived muscle after 12 days of differentiation. (scale bar = 1 mm). Higher magnification immunostaining images showing myosin-expressing fibers (MF20, pink) composed of multiple fused nuclei (NucB, blue) (scale bar = 200 µm). **b)** Representative traces of muscle contracilte force in response to 1 Hz electrical stimulation. Unpaired t-tests comparing the **c)** maximum contraction force (t(9) = 0.47, p = 0.65) and **d)** time-to-peak-force (t(9) = 3.59, p = 0.006) between WT (n=5) and RGECO C2C12 (n=6) muscles. **e)** Principal component analysis plot derived from RNAseq STAR-Salmon alignment and quantification data. **f)** Heatmap of normalized muscle-relevant gene expression for WT and RGECO C2C12 displayed on a log_2_ axis. **g)** Normalized gene expression for protein-coding genes of myosin isoforms, ordered by increasing maturity along the myogenic timeline. Two-way ANOVA (F(5,54) = 11.67, p < 0.0001) with post-hoc Šídák’s multiple comparisons tests was performed to find significant differences between groups (p<0.001***, p<0.0001****).

Electrical stimulation of engineered muscle tissues after 1 week in culture (1 Hz, 50 ms pulse width) showed that both WT and RGECO C2C12-derived muscle tissues were contractile in response to stimulation (Figure 4b). Videos of muscle displacement were quantified using our established open-source algorithms^27,33^ and converted into values of force using measured mechanical properties of the GelMA hydrogel substrate (Figure S3). While both cell types generated muscle tissues capable of similar maximum contractile forces (Figure 4c), RGECO C2C12-derived muscles demonstrated slightly altered contractile dynamics with a reduced time required to reach peak force (Figure 4d).

To further quantify differences between WT and RGECO C2C12-derived muscles, we conducted bulk RNA sequencing (RNAseq) of tissues fabricated within our Limb-on-a-Chip devices. While principal component analysis (PCA) showed separate clustering of WT and RGECO C2C12 (Figure 4e), heatmaps of genes that play key functional roles in the muscle contractile apparatus^34^ showed minimal differences in muscle-relevant gene expression between groups (Figure 4f). Importantly, despite differences in the expression of embryonic and postnatal myogenic markers such as myosin heavy chain 3 and 8 (Myh3, Myh8), muscle tissues derived from both WT and RGECO C2C12 demonstrated similar express of all mature myosin isoforms including Myh1, Myh2, Myh4, and Myh7 (Figure 4g). Taken together, these experiments validated the ability to fabricate aligned, mature, and contractile skeletal muscle tissues within our Limb-on-a-Chip platform using both WT C2C12 that enable tissue-wide force readouts and RGECO C2C12 that are compatible with both tissue-wide force readouts and single-fiber activation recording. As a result, RGECO C2C12 were used for future co-culture experiments with motor neurons.

### Compartmentalized Co-Culture of Motor Neurons and Muscle in Limb-on-a-Chip

Establishing relevance of our Limb-on-a-Chip platform for neuromuscular co-cultures required first validating motor neuron growth and differentiation within devices in monoculture, followed by demonstration of compartmentalized long-term co-cultures with muscle tissues. We leveraged established protocols^27,35^ to differentiate mouse embryonic stem cells (mESCs) into spheroids expressing a green fluorescent marker of Hb9 gene expression, indicating differentiation into a motor neuron lineage (Figure 5a). Individual spheroids suspended in collagen were seeded into separate motor neuron compartments within a Limb-on-a-Chip device for both monoculture and muscle co-culture experiments with RGECO C2C12 myoblasts (Figure 5b). Both monoculture and co-culture tissues were maintained in the same media, composed of a cocktail of muscle differentiation media 50% (v/v) and neuron differentiation media 50% (v/v) containing glial derived neurotrophic factor (GDNF) and ciliary neurotrophic factor (CNTF).

**Figure 5.**
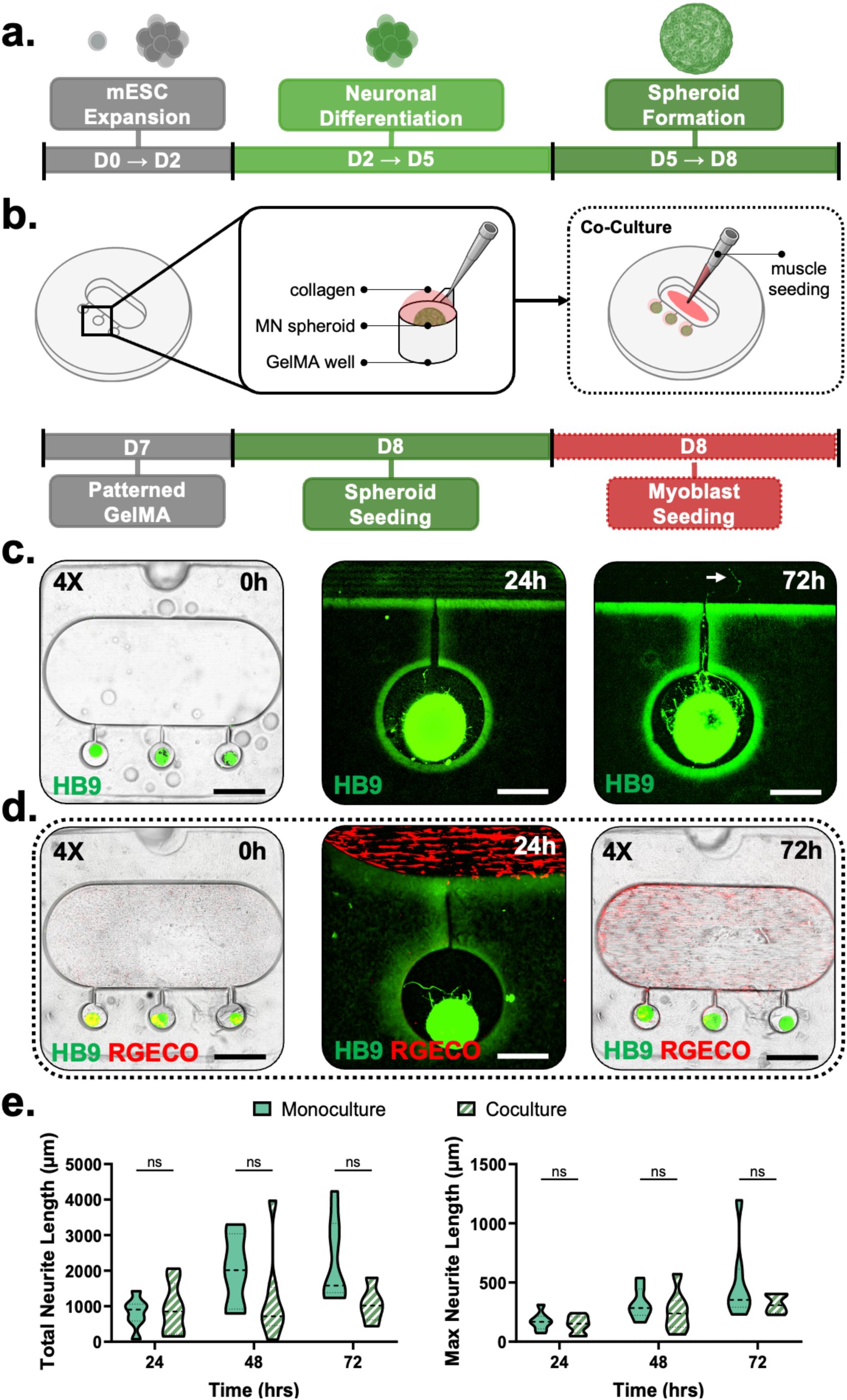
Compartmentalized Co-Culture of Motor Neurons and Muscle in Limb-on-a-Chip. **a)** Characterization of motor neuron differentiation timeline. **b)** Schematic of Limb-on-a-Chip seeding for both neuron mono-culture and neuromuscular co-culture experiments performed on day 8 of differentiation. **c)** Monoculture Experiment: (Left) Image of pseudo-brightfield overlaid with fluorescent HB9 signal. Progression of neurite outgrowth at 10X magnification for (Middle) 24 and (Right) 72 hours after Limb-on-a-Chip seeding. **d)** Co-culture Experiment: Images of co-cultures from the day we seeded (Left) to 24 hours after seeding (Middle) and 72 hours after seeding (Right). **e)** Quantification of neurite outgrowth of both monoculture (n=7) and co-culture (n=8, 24 h; n=7, 48 h; n=5, 72 h) wells using a two-way ANOVA for total neurite length (F(2,35) = 1.85, p = 0.173) and maximum neurite length (F(2,35) = 0.655, p = 0.526). Post-hoc Šídák’s multiple comparisons tests were performed to compare monoculture to co-culture at each timepoint. 4X images display black scale bars 2 mm in length. Higher magnification fluorescent images display 500 µm white scale bars.

Monoculture experiments showed that motor neuron spheroids were reproducibly placed within each compartment of the “spinal cord” (Figure 5c). Motor neurons projected axons that migrated throughout the compartment and grew into the micro-channel connecting the neuron and muscle chambers within 72 hours of culture. Similarly, co-cultures of motor neuron spheroids and myoblasts demonstrated clear spatial segregation and differentiation of both cell types in separate compartments (Figure 5d). To verify whether the presence of muscle influenced the early stages of neuronal growth, we quantified the total length of all axons projecting out of the motor neuron spheroid, as well as the length of the longest axon projection from the spheroid. Our experiments revealed no significant differences in axon outgrowth between monocultures and co-cultures within the first 72 hours of culture (Figure 5e).

Neuromuscular co-cultures within Limb-on-a-Chip devices were maintained over 10 days before recording videos of muscle activity as indicated by the genetically encoded calcium sensor (Figure 6a, Video S2). Videos were processed via an algorithm that used the maximum fluorescence intensity projection over time to segment and individually label all active muscle fibers within the tissue (Figure 6b). This image processing workflow enabled tracking the activity of several muscle fibers within the tissue with single cell precision (Figure 6c), highlighting the future potential of our Limb-on-a-Chip platform for tracking muscle innervation status in long-term neuromuscular co-cultures.

**Figure 6.**
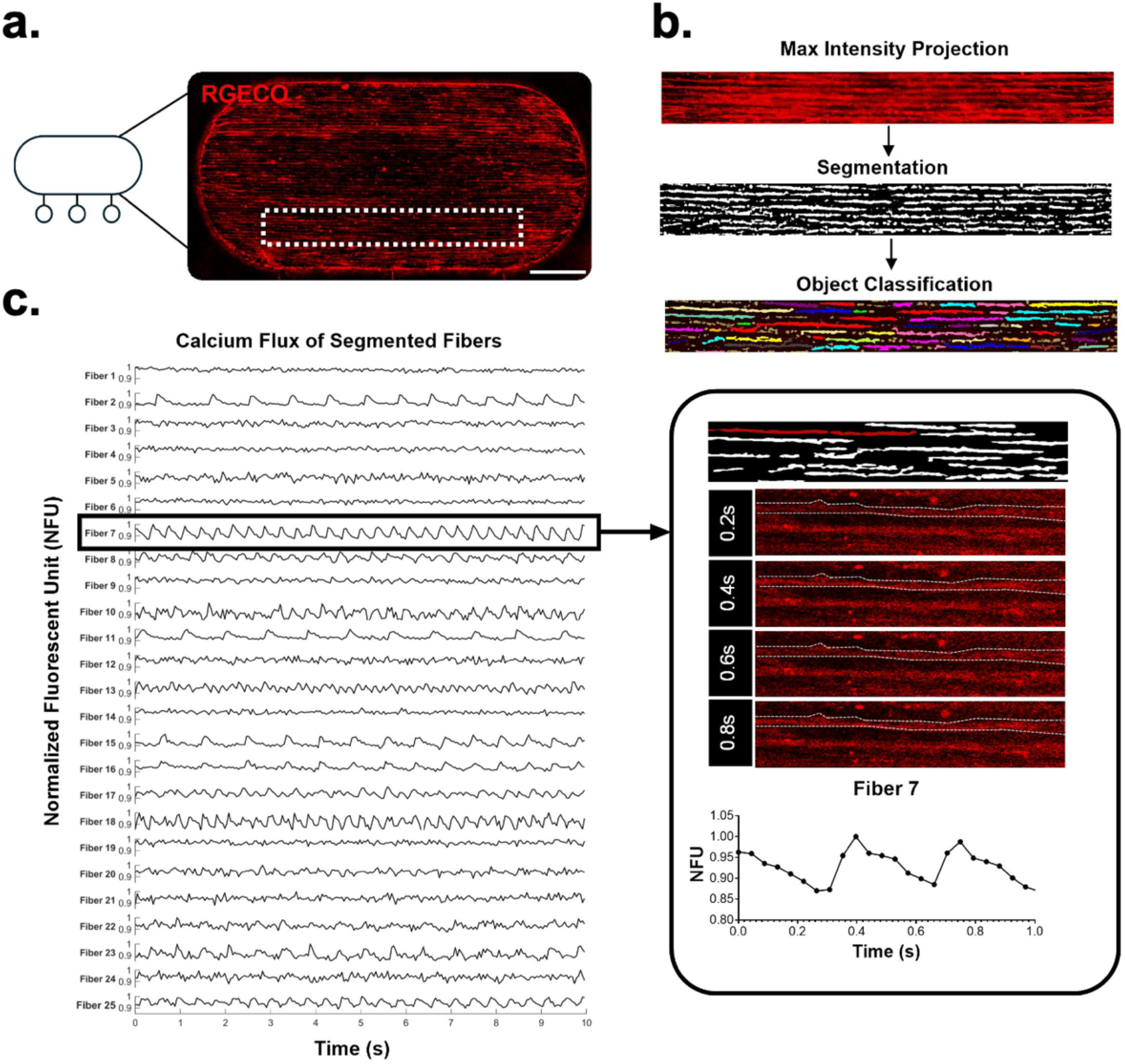
Monitoring Muscle Fiber Activation in Neuromuscular Co-Cultures. **a)** Muscle compartment of Limb-on-a-Chip after 10 days of co-culture with motor neurons. (scale bar = 1 mm). **b)** Data analysis pipeline used to analyze calcium flux of RGECO C2C12 muscle fibers. First, maximum intensity projections over time are created to identify all active muscle fibers within a video, serving as the input for subsequent calcium tracking steps. Then, an open-source random forest classifier (ilastik) is used to segment the maximum intensity projection, with black indicating background and white indicating an active muscle fiber. This mask was used as an input in the object classification step, where the mask of all active fibers ROI is applied to the raw video (10 seconds, 22.7 fps) in the object classification workflow in ilastik. All small, noisy “objects” are filtered out in this step, leaving only smooth fibers. Finally, the mean intensity for every object is generated for every frame of the raw video, creating a trace of calcium flux within each identified active muscle fiber. **c)** Calcium traces plotted for 25 out of 48 total identified muscle fibers in the imaging window. Fiber 7 is highlighted in the inset to showcase how fluorescence change in the raw videos is converted into the resultant calcium activity trace.

## DISCUSSION

Reproducible *in vitro* models of the neuromuscular interface that enable quantitative readouts of tissue activity have the potential to advance understanding of the neuromuscular system in health and disease.^26^ Existing model systems, ranging from organoids to organs-on-a-chip, have provided significant insight into neuromuscular biology.^5^ While self-assembled 3D organoids and 2D co-cultures enable forming functional neuromuscular junctions (that are more readily visualized in 2D cultures), they are typically not compatible with quantitative force readouts and rely on stochastic self-assembly processes that limit inter-tissue reproducibility. Organ-on-a-chip devices enable more repeatable fabrication, and often even separate motor neuron soma and muscle fibers to better mimic *in vivo* architecture. However, such compartmentalized devices typically rely on 3D tissues which, while compatible with whole-tissue force readouts, do not enable monitoring the activity of single muscle fibers. Taken together, these challenges emphasize the need for new platforms that address the limitations of current tools.

We have developed a Limb-on-a-Chip device that combines the advantages of compartmentalized cultures, as in 3D organs-on-a-chip, with the benefits of high-resolution imaging compatibility, as in 2D co-cultures. Our platform leverages a 1-step molding method to pattern an all-hydrogel chip composed of a “spinal cord” housing independent motor neuron spheroids connected via micro-channels to a downstream muscle “limb”. This format enables reproducible and compartmentalized co-cultures of motor neurons and skeletal muscle, and is compatible with functional tissue-wide force readouts, similar to 3D organs-on-a-chip. Moreover, because our Limb-on-a-Chip maintains a 2D morphology at the neuromuscular interface, it also enables single-cell resolution in monitoring muscle fiber activity, similar to 2D self-assembled co-cultures. Finally, the open-well format of our device enables culturing engineered neuromuscular tissues in the same media, ensuring biochemical crosstalk between cell types that has been shown to be beneficial.^27^

In this study, we showcase the ability to accurately fabricate Limb-on-a-Chip devices with precisely defined and long-lasting microscale features (Figures 2-3), and also demonstrate that our devices enable compartmentalized monocultures of muscles or motor neurons, as well as compartmentalized co-cultures of muscles and motor neurons (Figures 4-5). Furthermore, we showcase the ability to quantitative monitor tissue-wide force (Figure 4) and single fiber calcium activity (Figure 6). An advantage of our method is that the 3D printed STAMP molds can be readily modified to modify chip features, such as the number and size of motor neuron and muscle compartments, or the architecture of the micro-channels that connect the two cell types. A limitation of our study is that we did not screen a wide range of hydrogel formulations for fabricating Limb-on-a-Chip devices, highlighting opportunities to identify substrates other than 7.5% GelMA that better promote muscle and motor neuron differentiation while still enabling compartmentalized co-cultures. However, given the compatibility of our STAMP approach with a range of cross-linkable hydrogels, we anticipate that such optimization can be readily performed and adapted to the needs of specific cell lines. Another limitation of our study is that we used mouse cell lines to derive motor neurons and muscles, which do not fully represent human biology.^1^ Our previous studies have demonstrated that STAMP is also compatible with human cells,^16^ highlighting an opportunity to extend the Limb-on-a-Chip platform to human neuromuscular co-cultures in the future.

Our goal with this study was to showcase the ability to accomplish compartmentalized neuromuscular co-cultures compatible with functional readouts at the tissue- and single cell-scale. We anticipate that future studies can leverage the Limb-on-a-Chip system to advance biological understanding of neuromuscular assembly, maturation, and adaptation in both physiological and pathological conditions. For example, while we did not observe significant differences in axonal outgrowth between monocultures and co-cultures within the first 72 hours of cell seeding, future studies that investigate differences in the rate and magnitude of axonal outgrowth over longer-term cultures may lend further biological insight into how crosstalk between cells promotes neuromuscular interface development.

In summary, we have developed an accessible protocol that enables researchers to fabricate reproducible, compartmentalized, and scalable models of the neuromuscular interface. We anticipate our protocol’s use of 3D printed molds could allow others to make custom modifications to Limb-on-a-Chip architecture as needed, such as changes in micro-patterned feature sizes, or the addition of new chambers to enable co-cultures with other cell types. By advancing the ability to reliably fabricate compartmentalized *in vitro* neuromuscular models compatible with quantitative high-resolution functional readouts, we hope that our Limb-on-a-Chip advances the state-of-the-art in modeling the neuromuscular system.

## METHODS

### Device Fabrication

#### STAMP Fabrication

Limb-on-a-Chip STAMPs (Figure 2a-b) were designed using CAD software (Solidworks) and prepared as previously described.^16^ Briefly, the portion of the STAMP containing the main microfeatures of the Limb-on-a-Chip was printed in UpPhoto resin with the NanoOne 2-photon printer (UpNano GmbH) with a 10x objective after slicing exported .stls with UpNano’s slicer software Think3D (UpNano GmbH). STAMPs were post-processed in 3 subsequent isopropyl alcohol baths for 10 minutes each and then allowed to further photo-cure for at least 12 hours before assembly to holders printed using stereolithography (SLA, see Figure 2d). SLA printed holders were designed in CAD software (Solidworks) and then sliced with Preform slicing software (Formlabs) and printed using a Formlabs 3B Stereolithography printer with Biomed Clear Resin (Formlabs). The holders were post-processed in isopropyl alcohol in the Form Wash for 30 minutes and then cured in the Form Cure for 60 minutes at 60 °C. The holders were then run through a Tuttnauer benchtop line autoclave (Tuttnauer USA Co Ltd, 2840EL-D) at 121 °C using the preset plastic cycle and allowed to cool to room temperature. Subsequently, the micropatterned UpPhoto stamps were adhered to the biomed clear holders with a thin layer of superglue (J-B Weld, JB-JBW-33120) to create the whole device.

#### GelMA Solution

7.5% w/v Gelatin Methacrylate (GelMA) solution with 0.5% w/v Lithium phenyl-2,4,6-trimethylbenzoylphosphinate (LAP) (900889, Sigma-Aldrich) was produced by first reconstituting 0.5% w/v LAP in DMEM (D6429-500ML, Sigma Aldrich) at 37 °C for 30 minutes until fully dissolved. After filtering the LAP solution, 7.5% w/v Bovine 225 bloom GelMA (Cytoink) was reconstituted in the LAP solution at 37 °C until fully dissolved. The 7.5% w/v GelMA solution was stored at 4°C until further use.

#### Limb-On-A-Chip Micropatterning

STAMPs were sterilized via immersion in 3% H₂O₂ (FisherScientific) overnight at room temperature. Residual H₂O₂ was rinsed off the STAMPs by immersion in a PBS bath for at least 30 minutes at room temperature. STAMPs were then coated with 10% bovine serum albumin (BSA) (Thermo Fisher) for at least 1 hour at 37°C. We prepared the GelMA for micropatterning by warming it to 37°C. Glass-bottom 24-well plates (Cellvis) were plasma treated for 30 seconds per well using a handheld oxygen plasma treater to promote forming a flat layer of gel.

Once GelMA was sufficiently warmed and in a liquid state, we pipetted 350 μL of 7.5% GelMA into a plasma-treated well. We subsequently aspirated excess BSA from the stamps carefully and placed the Limb-on-a-Chip stamps into the filled wells. After ensuring there were no visible bubbles in the gel before polymerization, we photocrosslinked the GelMA around the Limb-on-a-chip stamp by resting the plate on top of a UV lamp for 2 minutes at approximately 20 mW power (Thorlabs). Then, the STAMPs were carefully removed, and each micropatterned gel comprising a Limb-on-a-Chip was washed once with PBS and kept hydrated in cell culture media.

To ensure Limb-on-a-Chip STAMPs could be reused with high feature fidelity, a three-part cleaning step was followed. First, STAMPs were soaked in a 0.1 mg/mL Liberase solution (Sigma Aldrich) for at least 1 hour at 37°C to degrade any residual GelMA stuck to the small features of the STAMP. Then, STAMPs were re-sterilized and cleaned using H₂O₂ for 30 minutes at room temperature, followed by a final PBS soak for an additional 30 minutes at room temperature. After cleaning, STAMPs could be coated with BSA, enabling reuse.

### Rheology

Rheological measurements were performed using a TA Instruments Discovery HR-2 hybrid rheometer equipped with a 20 mm parallel-plate geometry. All tests were conducted at 25 °C using a solvent trap to minimize water evaporation, with a gap distance between 1.0 and 1.3 mm. Amplitude sweep tests were performed at an angular frequency of 10 rad s⁻¹ over a strain range of 10⁻² to 10² % to determine the linear viscoelastic (LVE) region. Subsequently, frequency sweep tests were carried out within the LVE range at a fixed strain of 0.5 %, over an angular frequency range of 10⁻¹ to 10² rad s⁻¹.

### Cell Culture

#### Muscle Culture

C2C12 myoblasts were initially cultured on plastic tissue culture treated flasks using muscle growth medium which contained 10% fetal bovine serum (A5670701, Gibco), 1% penicillin-streptomycin (MT30002CI, Fisher Scientific), and 1% L-glutamine (MT25005CI, Fisher Scientific) in a high glucose formulation of DMEM with 4.5 g/L of glucose and sodium pyruvate (MT10013CV, Fisher Scientific). Myoblasts were passaged prior to reaching 70% confluence. C2C12s were prepared at a concentration of 3x10⁶ cells/mL cells/mL and seeded into the Limb-on-a-Chip muscle chamber using 10 μL of cell solution added to 5 μL of co-culture media for a total of 30k cells suspended in a total of 15 μL in the muscle chamber. Myoblasts were maintained in the chip in growth medium containing 1 mg/mL of aminocaproic acid (A2504, Sigma Aldrich). Upon reaching confluence in the chips, myoblasts were switched to differentiation medium to induce myoblast fusion and maturation. Muscle differentiation media consisted of 10% horse serum (26050-088, Gibco), 1% penicillin-streptomycin, 1% L-glutamine, and 50 ng/mL of human insulin-like growth factor-1(I1146, Sigma-Aldrich) in a high glucose formulation of DMEM. Media was changed daily throughout the course of the experiments.

#### Neuronal Culture

Prior to beginning neuronal culture, we seeded gamma-irradiated mouse embryonic fibroblasts onto a 6-well plate to serve as a feeder layer for our mouse embryonic stem cell (mESC) culture. The following day, we seeded mESC onto the feeder layer using ES-LIF media, made up of EmbryoMax ES DMEM (SLM-220, Sigma-Aldrich) with 15% fetal bovine serum, 1% penicillin–streptomycin, 1% L-glutamine, 1% Embryo-Max nucleosides (ES-008, Sigma-Aldrich), 1% MEM non-essential amino acids (11140050, Thermofisher Scientific), 0.1 mM 𝛽-mercaptoethanol(21985023, Thermofisher), and 0.1% v/v ESGRO mLIF (26050070, Fisher Scientific). After 48 hours, we added our base neuronal medium (NDM) for 1 hour and then trypsinized the wellplate, seeding onto a tissue culture-treated 10 cm dish. Our NDM consisted of an initial half-and-half mixture of Advanced DMEM/F12(12634028, Thermofisher) and Neurobasal medium (21103049, Thermofisher) with 10% KnockOut Serum Replacement (10828010, Thermofisher), 1% pencillin-streptomycin, 1% L-glutamine, and 0.1 mM of 𝛽-mercaptoethanol. After 24 hours, we collected the unadhered neuronal cells and moved them to a low adhesion 10 cm petri dish. On day 3 of neuronal differentiation, we collected the medium and spun it down to aspirate the base neuronal medium, adding fresh NDM media which then contained 1 μM of retinoic acid (R2625, Sigma-Aldrich) and purmorphamine (540220, Sigma-Aldrich) (NDM+). We observed the culture and added 3 mL of the neuronal formulation on day 4 of neuronal differentiation. On day 5, we collect the spheroids and refresh the medium, now containing 10 ng/mL of ciliary neurotrophic factor(C3710, Sigma-Aldrich) and glial-derived neurotrophic factors(PR27022, Neuromics) (NDM++).

### Limb-On-A-Chip Seeding

#### Neuron Monoculture Seeding

Mature neuron spheroids (Day 8 in NDM++) were seeded in 2.4mg/mL porcine collagen (63800781, FujiFilm Scientific Inc.). The collagen was crosslinked using an 8:1:1 ratio of parts A, B, and C respectively, all mixed over ice and using chilled pipette tips. Specifically, spheroids for seeding were aliquoted in an Eppendorf tube and allowed to settle to the bottom. Any supernatant was carefully removed, and spheroids were resuspended in 30 μL of collagen. Then, spheroids were seeded into the three neuron wells of each Limb-on-a-chip suspended in collagen. This seeding was followed by a 30-minute gelation step at 37 °C to ensure that the collagen fully crosslinked. Then, the whole chip was flooded with 150 μL of collagen and set aside for an additional 30 minutes in the incubator to fully cross-link. This allows for a homogeneous coating of the Limb-on-a-chip and provides a favorable microenvironment to assess neural growth on this micropatterned substrate.

#### Neuromuscular Co-culture Seeding

On day 8 of differentiation, neuronal spheroids are resuspended in pure collagen and pipetted into the microwells generated by STAMP micropatterning. Approximately 2 μL of collagen containing one spheroid were deposited into each microwell, and all microwells within a Limb-on-a-Chip were seeded and allowed to cross-link, similar to the monoculture chip seeding. To ensure that the muscle chamber is free of residual media, we utilized a gel loading pipette tip to aspirate any remaining liquid. Next, we pipetted 5 μL of co-culture media, consisting of 50% muscle differentiation medium and 50% NDM++, into the muscle chamber. Then, the C2C12s were prepared at a concentration of 3x10⁶ cells/mL and seeded into the chamber using 10 μL of cell solution added to 5 μL of co-culture media for a total of 30k cells suspended in a total of 15 μL in the muscle chamber. The initial 5 μL was used to promote a homogenous distribution of cells in the chamber. The amount of media utilized was optimized to prevent fluid flow into the neuron wells. After 45 minutes, once the myoblasts have adhered, we proceeded to aspirate the media from the chip. We then flooded it with 150μL of collagen and set the chip aside for another 30 minutes to cross-link. Finally, we add co-culture medium into the chips after the 30-minute cross-linking period.

### Muscle Electrical Stimulation

Electrical stimulation was performed 14 days after being switched to differentiation media during muscle-only characterization studies. Cultures were placed in 2 mL of DMEM and supplied with 10 Vpp 5% duty cycle square waves at a frequency of 1 Hz using a Siglent SDG1032X function generator connected to electrodes made of platinum wires (Thermo Fisher Scientific, 010286.BY) suspended approximately 7 mm apart and 9 mm above the culture. Muscle twitch videos were recorded in brightfield with a 10x objective at 30 fps with a Zeiss Primovert Microscope.

### Bulk RNA Sequencing

RNA extraction was performed on day 14 of muscle differentiation using the Qiagen(74104, Hilden, Germany) RNeasy Mini Kit following the manufacturer’s instructions. The muscle monolayers were treated with the lysis buffer for 15 minutes before being spun down in QIAshredder spin columns(79656, Qiagen) to homogenize the tissue. Sequencing libraries were generated using the NEBNext® Ultra™ II Directional RNA preparation with poly(A) selection kit. Library sizes were quantified and verified via qPCR and Fragment Analyzer before being loaded on the Singular G4 platform in a 50-base paired-end configuration with 8+8 nucleotide indexes. The reads were then mapped against mm39 (GRCMm39) using the BWA-MEM algorithm. Quality control metrics such as mapping rates, unique 20-mers and fraction of ribosomal RNAs were calculated using bedtools version 2.30.0. FASTQ files were processed using the nf-core/rnaseq version 3.17.0 pipeline. We utilized the GRCm39 reference genome and ENSEMBL GRCm39 murine annotations. Differential gene expression was done in DESeq2 in R using the apeglm method(Zhu et al. (2018)) to provide Bayesian shrinkage estimators for effect sizes. We reported log_2_ fold changes and Benjamini-Hochberg adjusted p-values.

### Immunofluorescent Staining

Samples were fixed in 4% paraformaldehyde for 15 minutes then washed with PBS. Then, the samples were permeabilized using a 0.5% solution of Triton X-100(A16046.AE, Thermofisher) in PBS for another 15 minutes. Followed by an hour incubation in a blocking buffer of 10% BSA in PBS. For myosin-4 staining, the samples were left at 4C overnight with the blocking buffer previously mentioned with 0.5% Triton X-100, 1:200 dilution of conjugated anti-myosin 4(MF20) antibody to eFluor 660(50-6503-82, eBioscience), and 2 drops per mL of NucBlue Live ReadyProbes(R37605, Thermofisher). Finally, the samples were washed with PBS prior to imaging and submerged in Fluoromount aqueous mounting medium(Sigma-Aldrich, F4680).

### Image Acquisition

Daily fluorescence imaging was performed on a Nikon AXR confocal microscope, utilizing a 4X and 10X lens. Brightfield video recording of muscle contraction and stimulation was performed on a Zeiss inverted microscope. Stereoscope brightfield images of Limb-on-a-chip STAMP molds (figure 2e) were captured using a Leica Stereoscope.

### Data Analysis

#### Device Characterization

A Nikon AXR confocal microscope was used to image the autofluorescent regions of the GelMA Limb-on-a-Chip hydrogels created in this work and to characterize the dimensions of the grooves and microchannels of the hydrogel in comparison to modeled STAMP dimensions, as shown in Figure 3a–g. Groove z-stack images from multiple days within the same device were captured with a 20x objective lens over a range defined by the minimum and maximum heights of the grooves and then filtered with Nikon’s Denoise.AI feature. A cross-section of the z-stacks was then processed using ImageJ’s Orthogonal Views feature and subsequently segmented using the Segment Anything Model (SAM) (Meta AI) (see Figure 3a). Groove dimensions were then extracted by taking the boundary of the segmented image and using MATLAB’s findpeaks() function to identify groove boundaries by locating bounding valley regions (see Figure 3c). Average dimensions were calculated by aligning grooves at their first valley point and calculating average values at each location along the groove’s width. Heights were calculated as the distance between a groove’s first valley point and its peak value, while widths were calculated as the distance between two adjacent valley points.

Channel dimensions were characterized with 20x objective confocal images captured in the regions of the channels across multiple chips just after casting. After filtering the images with Nikon’s Denoise.AI feature, the images were further processed with a Gaussian filter to reduce peppering and then binarized with a global threshold of 0.5 using MATLAB’s imbinarize() function (see Figure 3f). 10 µm and 50 µm regions of the channels were cropped, and channel widths were calculated by skeletonizing each channel region and calculating the shortest distance to the edge of the segmented fiber from each skeleton point, using the MATLAB functions bwskel() and bwdist(). The final reported width for each channel section was then calculated by doubling all the measured distances from the edge of the binarized channel to the skeleton points and taking the average width value for each imaged channel (Figure 3g).

Leak-percentage characterization data were collected by seeding RGECO C2C12 myoblasts at a concentration of 3x10⁶ cells/mL in 10 µL into a well previously loaded with 5 µL of media. Reported leak percentages represent the number of channels where cells infiltrated divided by the total number of channels across all seeded devices.

#### Cell Culture Maturity Data Analysis

Neurite quantification was performed utilizing the daily immunofluorescent imaging of our HB9:GFP neurons. The max length and total neurite length were recorded for each sample and day using the ImageJ plugin NeuronJ.

#### Displacement Dynamics

Brightfield videos of muscle contraction were analyzed utilizing an open-source full-field displacement and strain tracking code from our prior published work.^33^ The code extracts spatial displacement maps and calculates mean average displacement values (MAD) metrics for each frame. We extracted max MAD values by taking the vertical distance between a valley and the subsequent peak as found by MATLAB’s findpeaks() function. Time to peak metrics were found by taking the time between an identified valley and the next adjacent identified peak.

### Statistical Analysis

Quantitative results are reported as mean ± standard deviation (SD). Details of sample sizes and statistical tests for each figure are included in the corresponding caption. For Figures 3d-e, One-way ANOVA and Kruskal-Wallis tests were performed using Python’s scipy.stats package. For Figure 3e, one-way t-tests were performed using MATLAB’s ttest() function, comparing designed CAD model values to measured mean widths. The remaining statistical tests were performed using Graphpad Prism 10. For muscle-only studies, when comparing metrics across both cell lines, a two-way ANOVA was performed with post-hoc Šídák’s multiple comparisons tests. Unpaired t-tests were used to compare contraction dynamics.

## Acknowledgements

The authors thank Prof. Rashid Bashir from the University of Illinois at Urbana-Champaign for the gift of muscle cells expressing genetically encoded calcium ion sensors that were used in this study. The authors acknowledge funding support from the DoD Office of Naval Research Young Investigator Program (N00014-24-1-2060, awarded to R.R.), DoD Army Research Office Early Career Program and PECASE (W911NF-22-1-0126,, awarded to R.R.), the NSF CAREER Program (2238715, awarded to R.R.), the DoD DURIP Program (W911NF-24-1-0106, awarded to R.R.), the NSF Graduate Research Fellowship Program (awarded to A.B.), the NIH T32 Neurobiological Engineering Training Program (awarded to L.S.), and the la Caixa Foundation Postdoctoral Fellowship (awarded to T.R.).

