## Supplemental Information for "Limb-on-a-Chip: An All-Hydrogel Platform for Scalable and Reproducible Engineering of Neuromuscular Tissues"

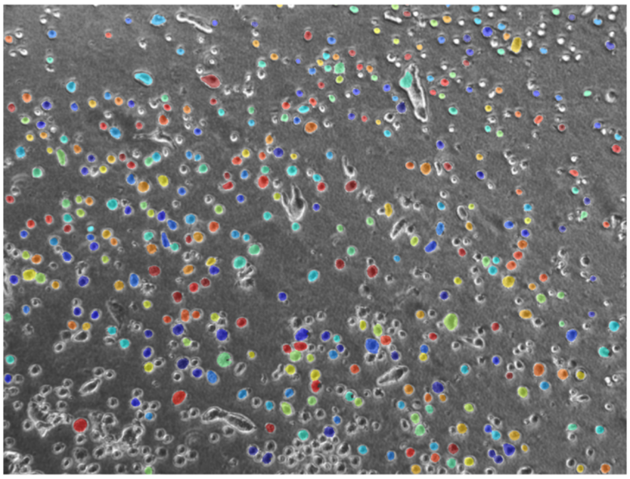

**Figure S1. Quantification of C2C12 diameter.** Example of Cellpose cell identification and labeling of a brightfield image of C2C12 WT myoblast cells in suspension. Colored regions denote different identified and labeled cell regions that were used to calculate equivalent diameters with the MATLAB regionprops() function.

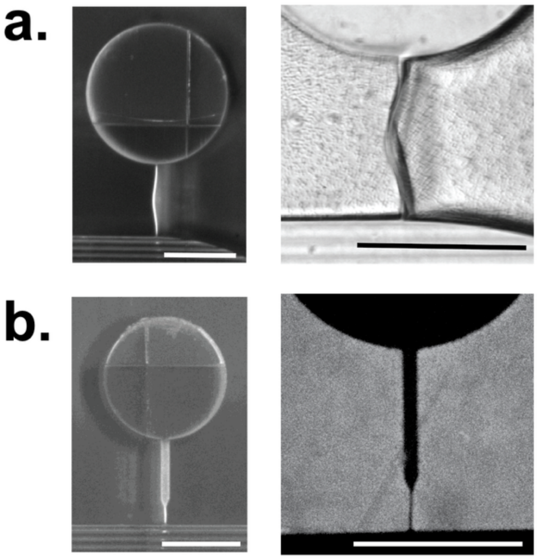


**Figure S2. Comparison of Different Channel Designs. a)** STAMPs with a straight 10 μm channel design (left) are used to mold GelMA hydrogels (right), highlighting the unstable nature of the thin channel that is prone to buckling (scale bars: 500 μm). **b)** Tapered channel design used in this study that more reliably patterned straight unwarped channels in both STAMPs (left) and STAMPed GelMA hydrogels (right) (scale bars: 500 μm).

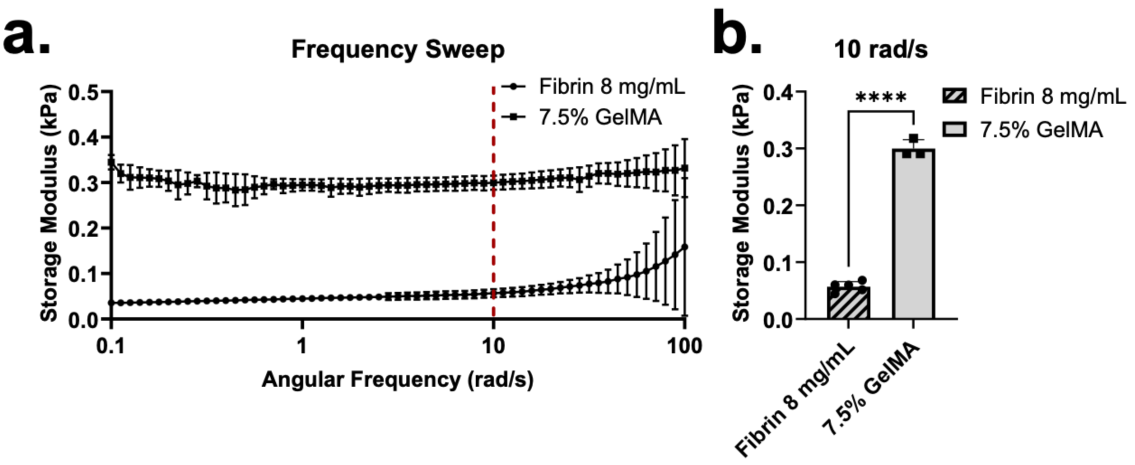


**Figure S3. Rheological Analysis of Hydrogels. a)** Frequency sweep of fibrin 8 mg/mL and 7.5% GelMA followed by **b)** a comparison of storage modulus at 10 rad/s. The storage modulus of the 7.5% GelMA hydrogel (0.30 ± 0.02 kPa, n = 3) was significantly higher than that of 8 mg/mL fibrin hydrogel (0.057 ± 0.009 kPa, n = 5). An unpaired t-test was used to compare a difference of means ($t(6) = 28.65$; $p<0.0001$****).

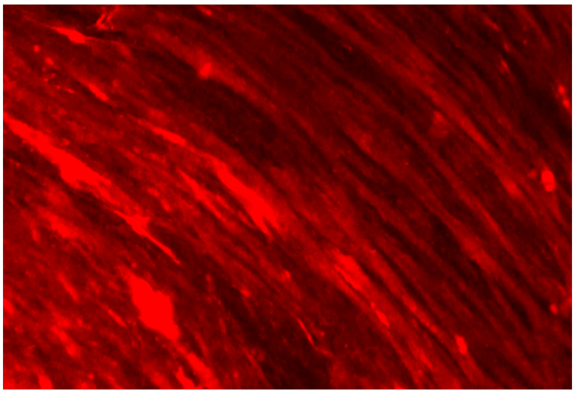

**Video S1.** Electrical stimulation (10 Vpp, 1Hz, 5% duty cycle) of RGECO C2C12 muscle on unpatterned hydrogel substrate showing both tissue-wide movement and single fiber-level calcium activity.

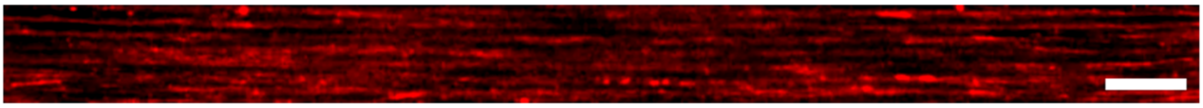

**Video S2.** Muscle calcium flux activity within Limb-on-a-Chip after 10 days of neuromuscular co-culture, recorded on a confocal microscope. (scale bar: 300 µm).
